# Novel Extended Tetraether Lipids Found in a High-CO_2_ Geyser

**DOI:** 10.64898/2026.01.06.697877

**Authors:** Janina Groninga, Leonie Wittig, Feriel Bouderka, Till L. V. Bornemann, Julius S. Lipp, Florence Schubotz, Saskia Keden, Alexander J. Probst, Kai-Uwe Hinrichs

**Author notes:** **Author details:** Leonie Wittig, Feriel Bouderka, Till L. V. Bornemann, Julius S. Lipp, Florence Schubotz, Saskia Keden, Alexander J. Probst, Kai-Uwe Hinrichs.

## Abstract

The growing research into the archaeal lipidome has uncovered a remarkable structural diversity in glycerol dialkyl glycerol tetraethers (GDGTs) and revealed complex membrane adaptations, especially in extreme environments. We performed a comprehensive analysis of the lipidome from the subsurface water of a CO_2_-rich, cold-water Geyser Andernach (Germany), using ultra-high-resolution mass spectrometry methods. We detected GDGT-0, presumably derived from the dominant community member *Candidatus* Altiarchaeum, providing evidence for its ability to synthesize tetraethers as previously predicted from metagenomic data. Beyond the typical GDGT-0 and acyclic glycerol trialkyl glycerol tetraether (GTGT-0), we discovered novel structural analogues, here referred to as extended GDGTs and GTGTs, characterized by the asymmetrical addition of up to two isoprenoid units to only one of their hydrocarbon side chains, analogous to those found in extended archaeols. The lack of GDGT ring synthase A (GrsA) and GrsB homologs in the corresponding metagenome-assembled genome, suggests that the producing archaeon may utilize extended GDGTs as a membrane adaptation to cope with the energy-depleted conditions of the geyser environment, highlighting the adaptive flexibility of archaea to extreme physicochemical conditions.

## 1 Introduction

Archaeal cell membranes differ fundamentally from those found in Bacteria or Eukaryotes, and harbor distinct biophysical properties that ensure cellular integrity even under hostile environmental conditions. This resilience is grounded in the molecular structure of their lipids, which comprise exclusively ether-linked isoprenoid chains attached to a glycerol-1-phosphate (G1P) backbone (Koga and Morii, 2007; Koga, 2012). Broadly, archaeal membrane lipids can be categorized into two main types: diether lipids, such as archaeols (ARs), which form conventional bilayer membranes, and tetraether lipids (glycerol dialkyl glycerol tetraethers; GDGTs), which build membrane-spanning monolayers (De Rosa and Gambacorta, 1988; Valentine, 2007).

Since the first description of GDGTs in the 1970s by Langworthy (Langworthy, 1977), the research on tetraether lipids has continued to grow, and over the subsequent decades, numerous structural modifications have been reported. These include the incorporation of up to eight cyclopentane rings (introduced via GDGT ring synthases (Grs) (Zeng et al., 2022)), or in addition to cyclopentane moieties one cyclohexane ring in the case of the *Nitrososphaerota*-specific crenarchaeol (Sinninghe Damsté et al., 2002). Other structural GDGT modifications include methylated derivatives in hyperthermophilic archaea (Knappy, 2010; Garcia et al., 2024), hydroxylated GDGTs, which have been proposed to act as temperature proxies (Liu et al., 2012a; Huguet et al., 2013, 2017), or unsaturated GDGTs (Zhu et al., 2014). Cross-linked GDGTs, known as glycerol monoalkyl glycerol tetraethers (GMGTs), in which the biphytanyl chains are connected via a covalent carbon-carbon bridge, represent another specific and unusual modification limited to thermophilic archaea (Morii et al., 1998; Knappy et al., 2011; Garcia et al., 2024). More recently, another unusual class of tetraether lipids, known as butanetriol dialkyl glycerol tetraethers (BDGTs) and pentanetriol glycerol tetraethers (PDGTs), which are characterized by a butanetriol or pentanetriol backbone, have been described (Becker et al., 2016; Coffinet et al., 2020). Initially, BDGTs were linked to the methanogen *Methanomassiliicoccus luminyensis* (Becker et al., 2016; Coffinet et al., 2020) but new evidence shows that Bathyarchaeia, a highly abundant archaeal class, are capable of BDGT synthesis as well (Dong et al., 2025). The diversity found in tetraether lipids not only reflects adaptations to various external conditions but also provides valuable chemotaxonomic indicators for different archaeal communities (Elling et al., 2015; Jensen et al., 2015; Feyhl-Buska et al., 2016; Elling et al., 2017; Zhou et al., 2020).

Similar modifications have been reported for AR, the diether precursors of membrane-spanning GDGTs (Lloyd et al., 2022; Zeng et al., 2022). Besides the typical AR with two C_20_-chains (2,3-di-*O*-phytanyl-*sn*-glycerol), extended variants that include one or two C_25_-chains (2-*O*-sesterterpanyl-3-*O*-phytanyl-*sn*-glycerol or 2,3-di-*O*-sesterterpanyl-*sn*-glycerol) have been described as another class of archaeal lipids that form membranes suited for strong osmotic stress and high salinity- (De Rosa et al., 1982; Vandier et al., 2021). Traditionally, these extended ARs (ext-ARs) were attributed to halophilic archaea inhabiting various hypersaline environments (De Rosa et al., 1982; Teixidor et al., 1993; Dawson et al., 2012; Birgel et al., 2014); however, recent findings support a more widespread occurrence than previously assumed (Becker et al., 2016; Natalicchio et al., 2017; Vandier et al., 2021). For instance, ext-AR and its intact derivatives, such as diglycosidic (2G) ext-AR, were identified in Crystal Geyser, a CO_2_-driven, cold-water geyser in Utah, USA (Probst et al., 2020) and in biofilms from the sulfidic Mühlbacher Schwefelquelle (Regensburg, Germany) (Probst et al., 2014; Perras et al., 2015). Notably, these extended ARs lipids have been attributed to the anaerobic, and CO_2_-dependent archaeon *Candidatus* Altiarchaeum of the order *Ca.* Altiarchaeales (also referred to as clade Alti-1, Bird et al., 2016) within the phylum Altiarchaeota, which represents the most abundant population in these aforementioned subsurface environments (Henneberger et al., 2006; Probst et al., 2014, 2018; Bornemann et al., 2022). Similar geochemical conditions likely promote the dominance of *Ca.* Altiarchaeum at our study site, the cold-water Geyser Andernach (GA), Germany (Bornemann et al., 2020, 2022), whose lipid inventory thus far remains unexplored.

Low-energy environments, such as the deep subsurface of the GA (Bornemann et al., 2022), often drive unique adaptations of cellular membranes and thus provide an ideal setting for discovering novel or unusual lipid structures. In this study, we describe yet another novel modification of GDGTs, characterized by extended isoprenoidal side chains, analogous to those found in extended ARs. These unusual extended GDGTs not only expand the known diversity of archaeal lipids but also provide new insight into potential alternative membrane adaptation strategies likely employed by *Ca*. Altiarchaeum in a deep subsurface environment.

## 2 Material and methods

### 2.1 Study site and sampling

Subsurface water samples were collected from the world’s highest erupting cold-water geyser that is situated in the Middle Rhine Valley near Koblenz in western Germany (50.448588° N, 7.375355° E) and was first drilled in 1903 (Bräuer et al., 2013; Dittrich, 2019). The GA reaches a temperature of around 18 °C and has a total depth of 351.5 m, with the upper 83 m of the borehole annulus being sealed with cement to prevent any surface water contamination (Dittrich, 2019). The up to 60 m high eruptions are fueled by geological degassing of mainly CO_2_ of magmatic origin and traces of H_2_ and H_2_S beneath the Eifel region and can be controlled with mechanical shutters (Bräuer et al., 2013; Barros et al., 2020). The concentrations of dissolved CO_2_ and HCO_3_^-^ can reach up to 1500 mg L^-1^ and 5700 mg L^-1^, respectively (Bornemann et al., 2022). These high-CO_2_ subsurface waters support a mesophilic microbial population dominated by *Ca.* Altiarchaeum (Bornemann et al., 2022).

Samples for lipid analysis were retrieved by collecting biomass from subsurface water onto glass fiber filters with a pore size of 0.7 µm (Ø = 142 mm; Whatman, 1825142) in June 2025, using an installed tap on the side of the geyser to prevent surface contamination. The water was filled into sterile containers and afterwards pumped through teflon tubes for filtration. The filters were immediately stored on dry ice on-site and then moved to -18 °C in the laboratory until further processing. On each filter, biomass from ∼80 to 140 L of subsurface water was collected. Additional materials from the GA had been collected for Bornemann et al., (2022) and, in addition to the therein described metagenomic analyses were also processed for lipid analysis. For these samples, water was collected in sterile containers during the eruption and subsequently filtered onto 0.1 µm polytetrafluoroethylene (PTFE) filters (Ø = 142 mm; Merck Millipore, JVWP14225).

### 2.2 Sample preparation and extraction

Total lipid extracts (TLEs) from the filters were obtained using the modified Bligh & Dyer protocol (Sturt et al., 2004). According to the protocol, the lipids were extracted in four steps using monopotassium phosphate buffer (8.7 g KH_2_PO_4_ per L; pH 7.4) for the first two extraction steps and a trichloroacetic acid buffer (50 g L^-1^; pH 2) for the following two extraction steps. Following each extraction, the samples were centrifuged for 10 min at 1250 rpm, after which the supernatant was decanted into a separatory funnel. Equal volumes of dichloromethane (DCM) and MilliQ water were added to induce phase separation. The aqueous phase was extracted three times with DCM, after which the combined organic phases were collected and subsequently washed three times using MilliQ water. The collected organic phases were evaporated under a gentle stream of N_2_ and the dry TLEs were stored at -18 °C until further processing.

### 2.3 UHPLC-ESI-timsTOF-MS^2^

Archaeal lipids were analyzed using a Dionex Ultimate 3000 ultra-high-performance liquid chromatography (UHPLC) system (Thermo Fisher Scientific, Bremen, Germany) connected to a trapped ion mobility time-of-flight mass spectrometer (timsTOF HT-MS), equipped with a vacuum-insulated probe-heated electrospray ionization (VIP-HESI) ion source in positive mode (Bruker Daltonics, Bremen, Germany). A reversed phase (RP) Acquity UPLC BEH C_18_ column (1.7 µm; 2.1 x 150 mm, Waters Corporation, Eschborn, Germany) fitted with a pre-column was used for chromatographic separation. The solvent system consisted of eluent A (MeOH:H_2_O (85:15) + 0.04% (v:v) HCO_2_H and 0.1% (v:v) NH_3(aq)_) as well as eluent B (MeOH:IPA (1:1) + 0.04% (v:v) HCO_2_H and 0.1% (v:v) NH_3(aq)_ and was used after the following two methods:

#### Method A

Following a method by Wörmer et al., (2013), lipids were separated at a column temperature of 65 °C and with a flow rate of 0.4 mL min^-1^ using the following gradient: initial conditions were at 0% eluent B and 100% eluent A, increasing to 15% eluent B by 2 min, 85% eluent B by 20 min, and reaching 100% eluent B by 28 min. The column was re-equilibrated at 0% eluent B until 34 min.

#### Method B

The separation was performed using a method based on Hopmans et al., (2021) with a column temperature of 30 °C with a flow rate of 0.2 mL min^-1^ and following a modified gradient (Meyer et al., 2025): starting with 5% eluent B and 95% eluent A for 3 min, eluent B was increased to 60% by 12 min, followed by a further increase to 100% eluent B over the next 38 min (up to 50 min total). The gradient was maintained until 80 min, followed by a decrease to 5% eluent B over 1 min, which was then held until the end of the gradient at 90 min.

Samples were dissolved in MeOH:DCM (9:1, v:v). Each measurement was started with a loop injection of 20 µL of calibration mix (10 mM sodium formate cluster solution:Agilent ESI-L tuning mix (1:4, v:v), Agilent Technologies, Santa Clara, USA) to enable an internal calibration of mass and ion mobility accuracy. The mass accuracy was better than 1 ppm and mobility calibration <0.2%.

The ESI *m/z* scan ranged from 50 to 3000 with positive ionization mode, the drying gas flow was set at 8.0 L min^-1^, drying gas temperature at 230 °C, nebulizer gas pressure at 4.0 bar, sheath gas at 300 °C, and sheath gas flow at 3 L min^-1^. Capillary voltage was set to 4500 V and end plate offset to 500 V. MS/MS stepping was applied to improve detection in the low and high mass range. For MS/MS analysis, a mobility-dependent collision energy table with two collision energies that were combined to one scan was used in parallel accumulation serial fragmentation (PASEF) mode with two ramps and a target intensity of 60000 and active exclusion for 6 seconds. The inverse mobility 1/K_0_ scan ranged from 0.80 to 2.30 Vs cm^-2^ with 100 ms ramp time.

DataAnalysis 6.2 (Bruker Daltonics, Bremen, Germany) was used for the data evaluation of ESI-timsTOF and APCI-QTOF measurements. Lipids were identified based on their exact masses, retention times, and diagnostic MS^2^ fragmentation patterns (Table S1-2 and Fig. S1-3). Individual ion chromatograms were extracted with a mass window of ± 0.01 Da, and quantification was performed relative to an injection standard (2 ng GTGT-C_46_ per injection, Huguet et al., 2006) added to the extract prior to the measurement. To account for differences in ionization efficiency among the various lipid species, we calculated response factors based on calibration curves of standard solutions containing structurally similar lipids, including GDGT-0, monoglycosidic GDGT-0 (1G-GDGT-0), AR, and diglycosidic AR (2G-AR) (Table S2).

### 2.4 UHPLC-APCI-QTOF-MS^2^

Measurements with atmospheric-pressure chemical ionization (APCI) were performed using a Dionex Ultimate 3000RS ultra-high-performance liquid chromatography (UHPLC) system (Thermo Fisher Scientific, Bremen, Germany) coupled to a maXis plus ultra-high-resolution quadrupole time-of-flight mass spectrometer (QTOF-MS) in positive APCI mode (Bruker Daltonics, Bremen, Germany). Two normal-phase Acquity UPLC BEH Amide columns (1.7 µm; 2.1 x 150 mm, Waters Corporation, Eschborn, Germany) fitted with a pre-column were used following the method by Becker et al., (2013). The flow rate was kept at 0.5 mL min^-1^ with a column temperature of 50 °C. The following gradient was used for eluent A (*n*-hexane) and eluent B (*n*-hexane:2-propanol (9:1, v:v)):

Starting with 3% eluent B and 97% eluent A for the first 2 min, the eluent B was increased to 5% and used until 10 min total. Afterwards, eluent B was increased to 10% for 10 min (up to 20 min total) and then further increased to 20% for 15 min (up to 35 min total). 50% of eluent B was used for the next 10 min (up to 45 min total) followed by 100% eluent B for 6 min (up to 51 min total). The column was equilibrated at 3% eluent B for 9 min (up to 60 min total).

Samples were dissolved in *n*-hexane:2-propanol (99.5:0.5, v:v). Each measurement was terminated with the injection of a calibration mix (ESI-L tuning mix, Agilent Technologies, Santa Clara, USA) to allow mass calibration. Each mass spectrum was lock mass-calibrated with a lock mass compound (922.0098 Da), which was added in the ion source. The mass accuracy was typically better than 1 ppm.

The APCI source parameters were as follows: scan *m/z* range from 50 to 2000, drying gas flow 6.0 L min^-1^, drying gas temperature 180 °C, nebulizer gas pressure 2.0 bar and collision energy 6 eV.

### 2.5 Detection of GDGT synthesis marker genes in Andernach metagenomes and reconstruction of their evolutionary history

To constrain the potential producer of GDGTs and its extended tetraether derivates in Geyser Andernach, we screened the metagenome assembly for homology of key marker genes involved in tetraether and GDGT cycloalkyl ring biosynthesis. Reference sequences of tetraether lipid synthesis genes (*tes*) were obtained from Zeng et al., (2022), totaling 1601 sequences. For the GDGT ring synthesis genes (*grs*), we included the *grsA* and *grsB* sequences identified in Zeng et al., (2019), all other archaeal homologs of these genes reported in that same study, and additional homologous *grsA* sequences obtained from the InterPro database for a total of 61 sequences (Blum et al., 2025). These reference datasets were used as queries to search for homologous sequences in the Geyser Andernach metagenomic assemblies. The assemblies were generated, as described in Bornemann et al., (2022) from environmental samples analoguos to those used for lipid analysis. Homology searches were performed using DIAMOND BLASTp (Buchfink et al., 2015, v2.1.13.167) with a minimum 50% amino acid identity, 50% alignment coverage, and an E-value cutoff of 1e^-5^. When present, corresponding hits were searched in the metagenome-assembled genomes (MAGs) generated from the same dataset, previously reconstructed in Bornemann et al., (2022). The identified MAG taxonomy was assigned according to the Genome Taxonomy Database (GTDB, Release R10-RS226) (Parks et al., 2025).

To reconstruct the phylogeny of the *tes* genes recovered from the GA metagenome, reference sequences from Zeng et al., (2022) were included to provide context. Recovered sequences were aligned alongside these references using MAFFT (Katoh et al., 2002, v7.407), and the resulting alignment was trimmed using TrimAl (Capella-Gutiérrez et al., 2009, v1.5.rev0). Maximum-likelihood phylogenetic trees were reconstructed from the trimmed alignment using IQ-TREE with the LG+C20+G+F model (Nguyen et al., 2015, v 3.0.1) and the resulting trees were visualized and annotated using iTOL (Letunic and Bork, 2021).

## 3 Results

### 3.1 Archaeal lipid composition in the Geyser Andernach

The investigation of the GA lipidome revealed a major proportion of glycosidic archaeols with up to three glucose moieties (Fig. 1, Table S1-2). Within the total archaeal lipid pool, 2G-AR (35%), followed by 1G-AR (15%), stood out as the most abundant lipid species. Additionally, tetraether lipids, specifically GDGT-0 and 1G-GDGT-0, were detected in considerable proportions, accounting for roughly 34% of the archaeal lipidome (Fig. 1b). Furthermore, approximately 10% of the archaeal lipid inventory consisted of lipids featuring extended versions of the usual phytanyl and biphytanyl moieties. Ext-ARs with either a 1G or 2G head group dominate this fraction and together represent more than half of the extended archaeal lipid pool (Fig. 1b). Most remarkable was the detection of several unusual tetraether lipids, tentatively identified as extended or di-extended GDGTs and GTGTs, which appear to comprise additional isoprenoid units, analogous to those found in ext-AR. Interestingly, extended GTGT-0 (ext-GTGT-0; 1.8% of total lipid pool) and di-extended GTGT-0 (di-ext-GTGT-0; 0.8% of total lipid pool) are more abundant than their extended GDGT analogs, which together only account for 0.1% of the total lipid pool. This trend is also evident in their glycosidic forms, as 1G-ext-GTGT-0 and 1G-di-ext-GTGT-0 were detected, whereas intacts extended GDGT remained below the detection limit.

**Fig. 1.**
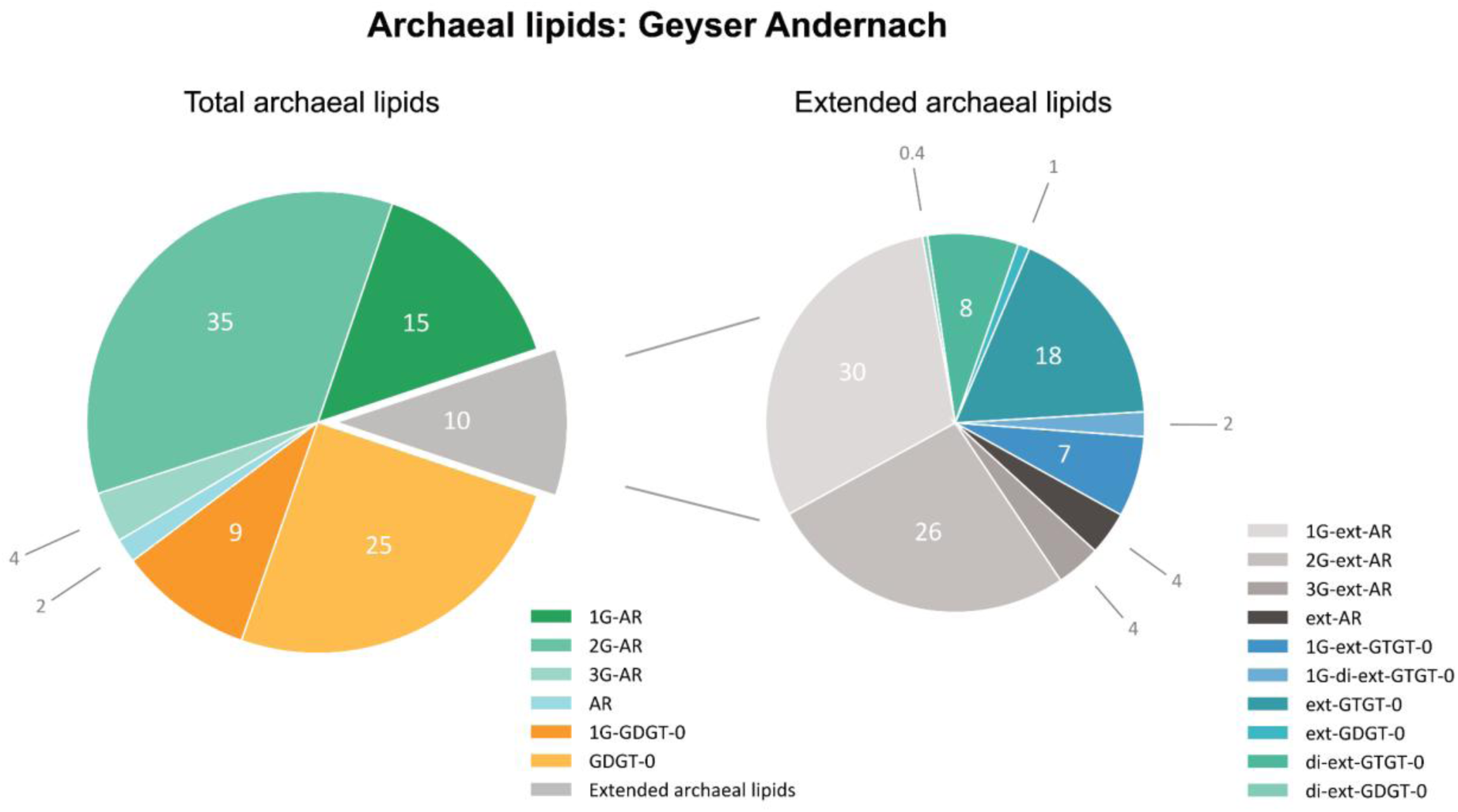
Relative abundances (%) of archaeal lipids detected in the erupting water from the Geyser Andernach (Germany), analyzed using reversed-phase UHPLC-ESI-timsTOF using method A. Corresponding lipid concentrations in ng/extract can be found in Table S1-2. The left chart shows the relative abundance of the total archaeal lipid inventory, with the grey sector showing lipids comprised of extended C_25_-sesterterpanyl chains. A breakdown of the extended archaeal lipid fraction, including novel compounds tentatively identified as extended GTGTs and extended GDGTs (c.f. 3.2), is featured in the right pie chart.

### 3.2 Structural characterization of extended GTGTs and GDGTs

Reversed phase UHPLC-ESI-timsTOF-MS revealed a distinct series of isoprenoidal tetraether lipids with retention times between 24 and 27 min, starting with GDGT-0 (24.4 min) and followed by two lipids tentatively annotated as ext-GTGT-0 (25.3 min) and di-ext-GTGT-0 (26.3 min) (Fig. **2**). In addition to the ∼1-min shift in retention time, the two extended compounds exhibited a stepwise mass increase, with a first increment of +72 Da relative to GDGT-0, reflecting one additional isoprenoid unit as well as two hydrogen atoms due to the tri-alkyl structure of GTGTs. The second increment of +70 Da corresponds to the addition of yet another isoprenoid unit within the GTGT lipid structure. Due to the close elution pattern of GDGTs and GTGTs, we performed additional normal-phase APCI-QTOF-MS^2^ analysis to facilitate better chromatographic separation and enable structural characterization based on the MS^2^ fragmentation pattern.

**Fig. 2.**
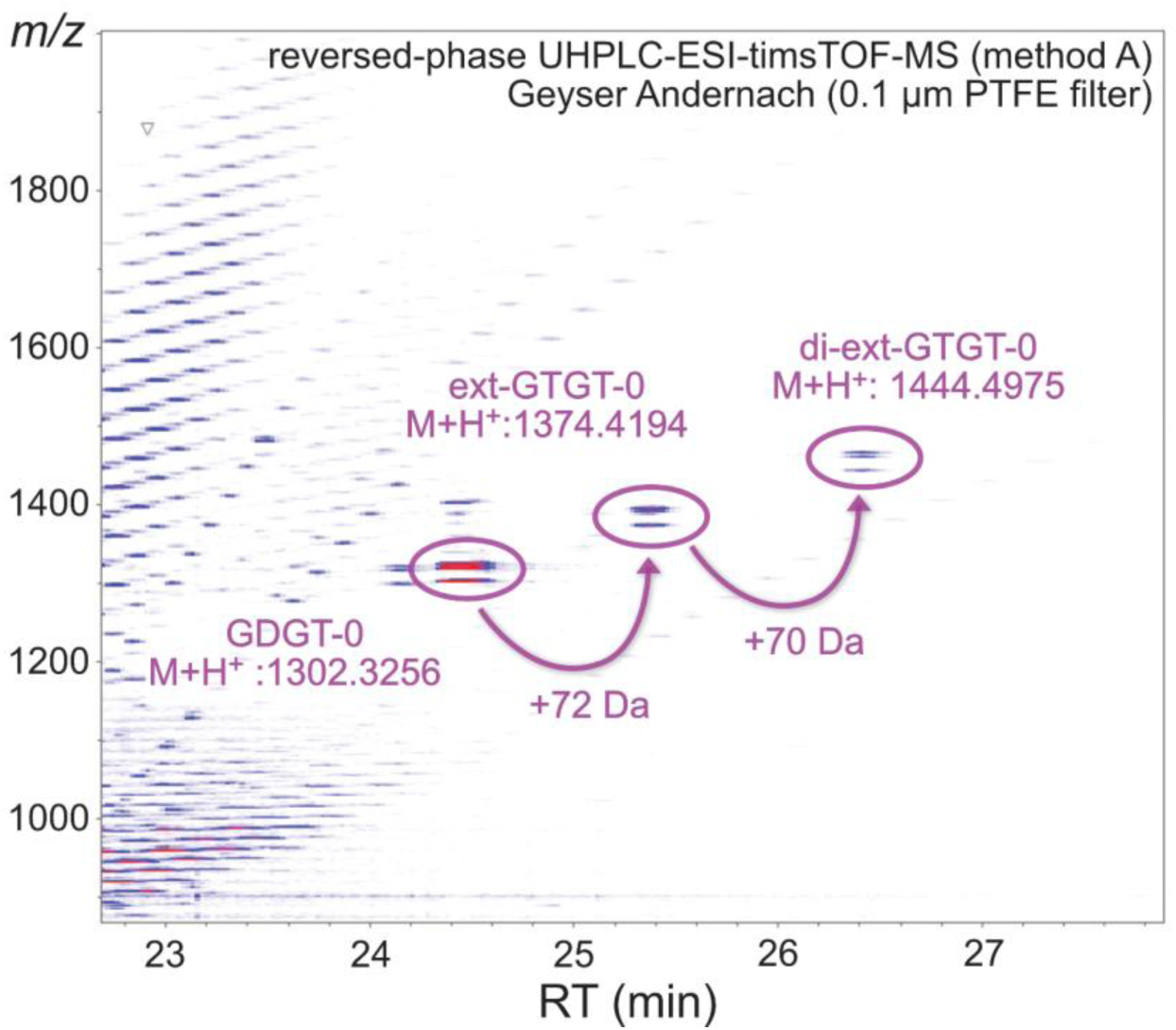
Heatmap of a positive RP-UHPLC-ESI-timsTOF-MS measurement (method A) of Andernach Geyser filter extracts highlighting GDGT-0, ext-GTGT-0, and di-ext-GTGT-0. Red color indicates higher signal intensities shown on a logarithmic scale.

UHPLC-APCI-QTOF-MS^2^ not only confirmed the distinct series of GTGTs but also revealed a corresponding GDGT series that was barely distinguishable in RP-UHPLC-ESI-timsTOF-MS. Starting with GTGT-0 ([C_86_H_174_O_6_+H]^+^: observed *m/z*: 1304.339; theoretical *m/z*: 1304.338) and GDGT-0 ([C_86_H_172_O_6_+H]^+^: observed *m/z*: 1302.326; theoretical *m/z*: 1302.323), respectively, each series is characterized by a mass increase of ∼70 Da (C_5_H_10_), consistent with the successive addition of isoprenoid units (Fig. 3). Next to the extended GDGTs and GTGTs, traces (<1.2% of the archaeal lipidome) of cycloalkylated GDGTs (GDGT-1 to GDGT-4) and glycerol monoalkyl glycerol tetraether (GMGT) were detected in the APCI-based measurements only (Table S3).

**Fig. 3.**
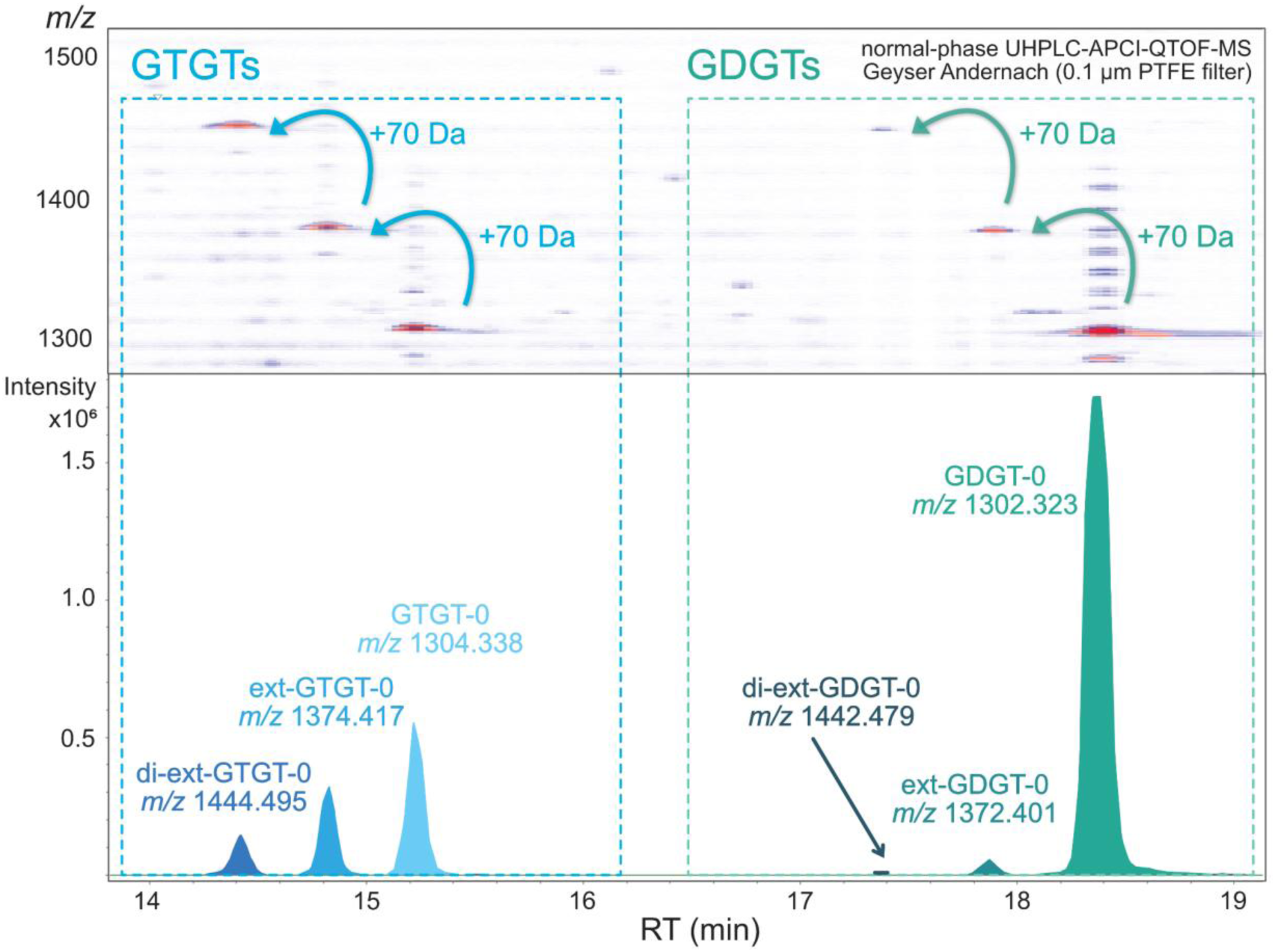
Heatmaps and extracted ion chromatograms illustrating the series of canonical and extended GTGTs and GDGTs in UHPLC-APCI-QTOF-MS analysis in extracts from the subsurface waters of the Geyser Andernach. Red coloring in the density map indicates high signal intensities shown in logarithmic scale.

UHPLC-APCI-MS^2^ spectra of *m/z* 1374.416 support the presence of one elongated side chain featuring an additional isoprenoid unit, as evidenced by a recurring mass increase of 70 Da throughout the MS^2^ spectrum when compared to the regular GTGT-0 (Fig. 4). Most prominent is the fragment ion at *m/z* 443.445 (C_28_H_58_O_3_) reflecting a C_25_-sesterterpanyl side chain attached to glycerol, alongside the regular C_20_-phytanyl chain attached to glycerol (*m/z* 373.367; C_23_H_48_O_3_) (Becker et al., 2016). The characteristic +70 Da isoprenoid extension can be traced across several higher-mass ions, most prominently at *m/z* 1002.055 ([M-C_23_H_48_O_3_+H]^+^) and *m/z* 1094.104 ([M-C_20_H_40_+H]^+^), reflecting the successive neutral loss of phytane (C_20_H_40_) and one glycerol unit (C_3_H_8_O_3_). The consistent occurrence of extended C_25_ chain fragments alongside those characteristic of the typical C_20_-phytanyl chain (e.g., *m/z* 373.368, *m/z* 931.978*, m/z* 949.987), and C_40_ chain fragments (e.g., *m/z* 743.712, *m/z* 651.665, *m/z* 615.644) support a general structure consistent with an extended GTGT-0 (Fig. 4 and Fig. S1).

**Fig. 4.**
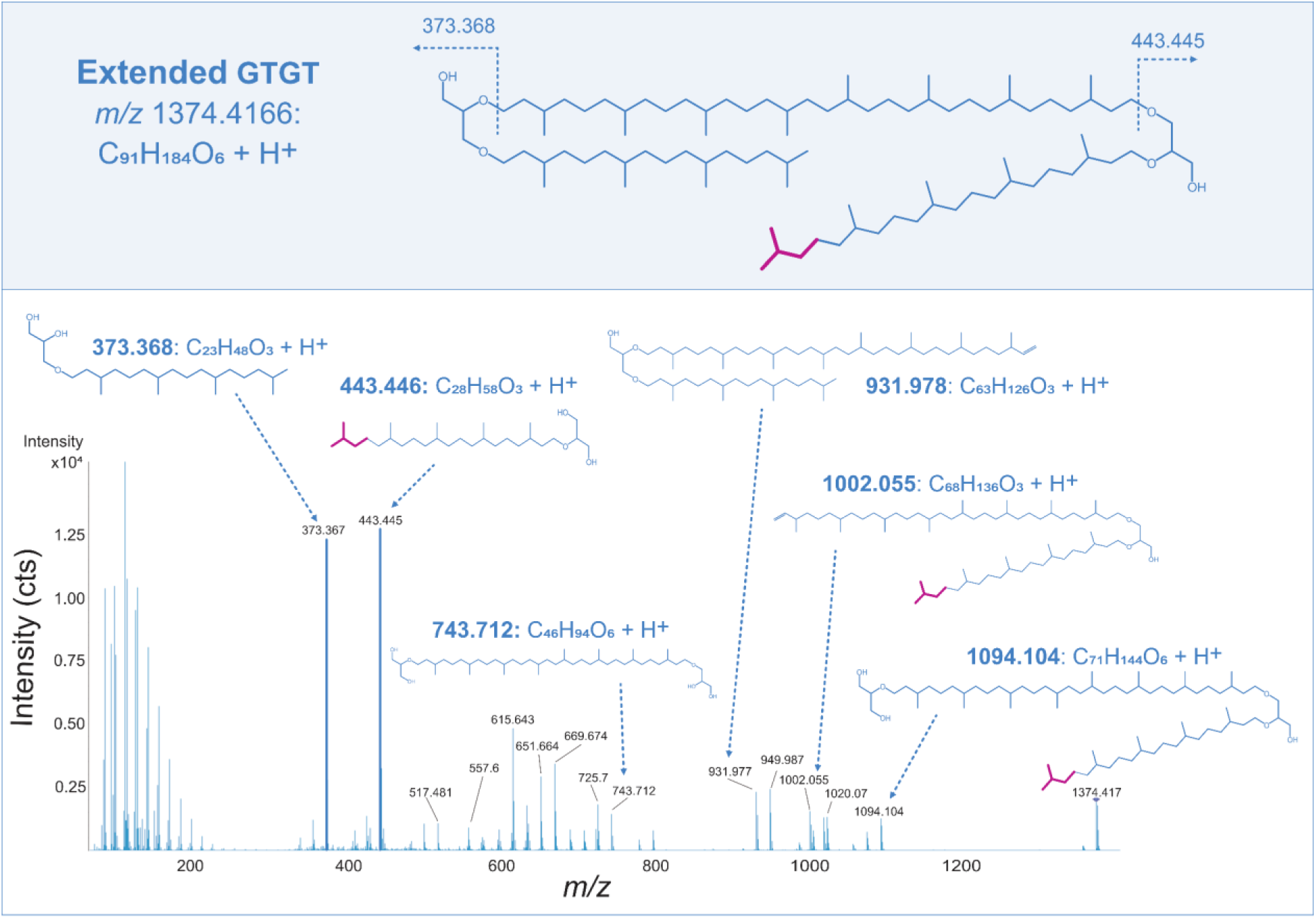
Proposed chemical structure of an extended GTGT-0 (C_91_H_184_O_6_) found in lipid extracts from the eruption waters of the Geyser Andernach (upper panel). UHPLC-APCI-QTOF-MS^2^ spectrum of extended GTGT-0 [M+H]^+^, emphasizing diagnostic MS^2^ fragment ions (lower panel). Magenta coloring indicates the additional isoprenoid unit characteristic for the extended GTGT-0 structure, which can be traced throughout the MS^2^ fragmentation pattern.

The earliest eluting compound of this series with *m/z* 1444.495 is consistent with the molecular formula [C_96_H_194_O_6_+H]^+^ (theoretical *m/z* 1444.495). Relative to GTGT-0 (C_86_H_174_O_6_; *m/z* 1304.338), this represents a mass increase of 140.156 Da, indicating the incorporation of two additional isoprenoid units (C_10_H_20_). The MS^2^ spectrum has the glycerol-C_25_-sesterterpanyl fragment (*m/z* 443.445) as base peak and lacks the characteristic glycerol-C_20_-phytanyl fragment ion at *m/z* 373.367. Fragment ions indicative of a C_40_ chain (e.g. *m/z* 743.712, *m/z* 651.665, *m/z* 615.644), and high intensity peaks at *m/z* 1094.105, *m/z* 1002.067, and *m/z* 443.445 corroborate a GTGT-0 with two extended C_25_ side chains (di-ext-GTGT-0) (Fig. 5 and Fig. S2).

**Fig. 5.**
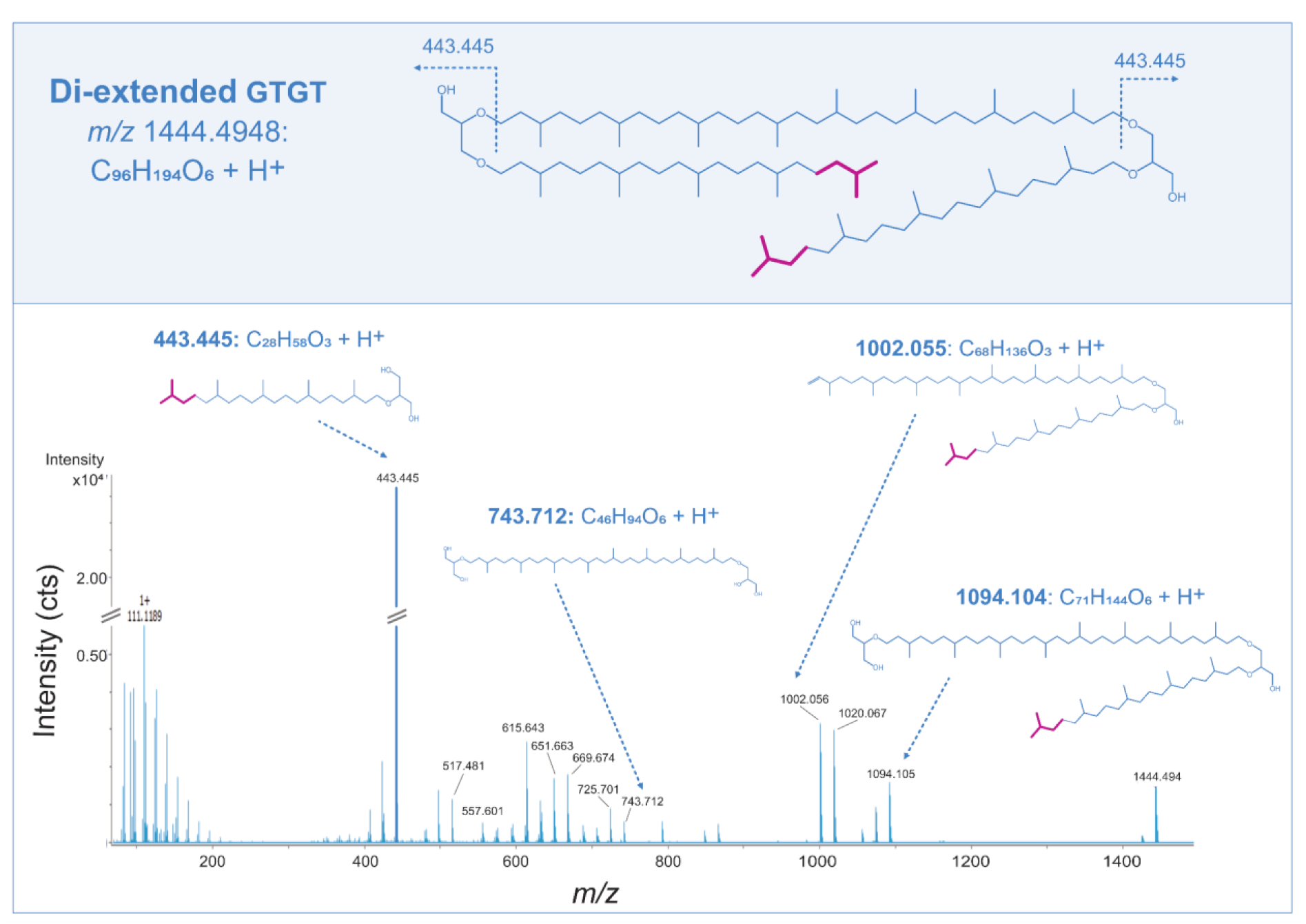
Proposed chemical structure of di-extended GTGT-0 (C_96_H_194_O_6_) found in lipid extracts from the eruption waters of the Geyser Andernach (upper panel). UHPLC-APCI-QTOF-MS^2^ spectrum of di-extended GTGT-0 [M+H]^+^ with characteristic MS^2^ fragment ions (lower panel). Magenta coloring indicates two additional isoprenoid units within the extended GTGT-0 structure.

Eluting roughly three minutes later, we observed an additional series of compounds, encompassing *m/z* 1302.324, corresponding to GDGT-0, *m/z* 1372.406, and *m/z* 1442.488, featuring the same successive addition of a +70 Da isoprenoid motif indicative of the novel extended GDGTs.

MS^2^ spectra of *m/z* 1372.406 ([C_91_H_182_O_6_ + H]^+^; theoretical *m/z* 1372.401) confirmed this hypothesis as we observed a high intensity fragment ion at *m/z* 813.792 reflecting a C_45_ side chain linked at both ends to a glycerol moiety, resulting from the loss of one C_40_-biphytanyl chain ([M-C_40_H_78_+H]^+^) (Fig. 6 & Fig. S3). Congruently, the fragment ions *m/z* 743.713, *m/z* 651.664 and *m/z* 615.642 typically observed in the GDGT core structure results from the neutral loss of one C_45_-nonaprenyl chain ([M-C_45_H_88_+H]^+^). This distinct fragmentation pattern is evidence of the presence of asymmetrical GDGTs that harbor one extended side chain. The distinct elution pattern in APCI-QTOF-MS and +70 Da mass increase alluded to additional occurrence of a di-extended GDGT presumably comprising one C_50_-decaprenyl chain. Due to the low abundance, no MS^2^ spectrum was available to further support the tentative structural assignment; however, we interpret *m/z* 1444.445 to contain one C_40_-biphytanyl chain and one extended C_50_-decaprenyl chain as opposed to two C_45_-nonaprenyl chains, due to the structural trend observed in the di-extended GTGT, where the C_40_-biphytanyl chain has been conserved (Fig. 5).

**Fig. 6.**
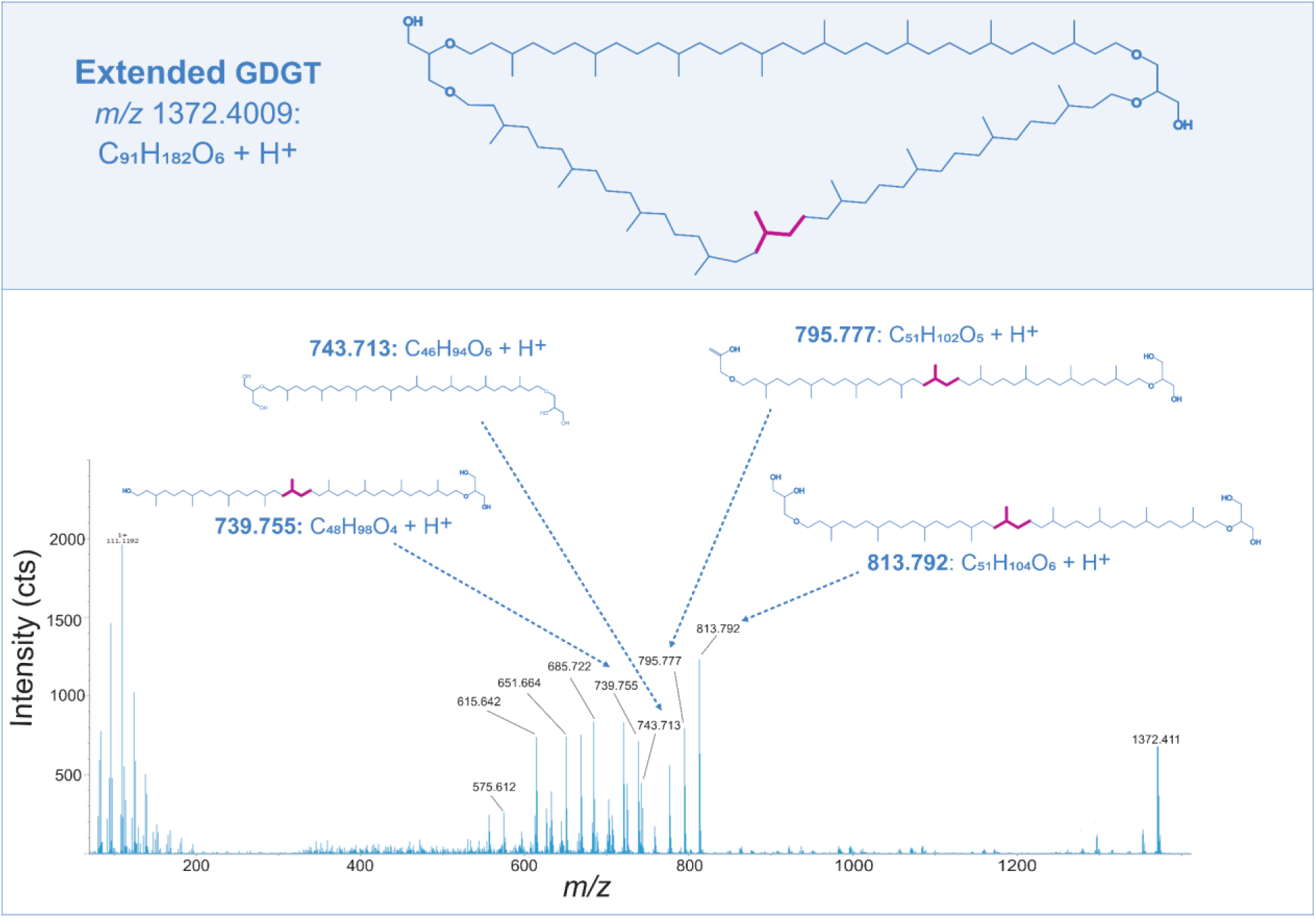
Proposed chemical structure of extended GDGT-0 (C_91_H_182_O_6_) found in lipid extracts from the eruption waters of the Geyser Andernach (upper panel). UHPLC-APCI-QTOF-MS^2^ spectrum of extended GDGT-0 [M+H]^+^, emphasizing diagnostic MS^2^ fragment ions (lower panel). Magenta coloring highlights the additional isoprenoid unit characteristic of the extended GDGT-0 structure, which is found throughout the MS^2^ fragmentation pattern.

Additionally, RP-timsTOF-MS^2^ measurements of glass fiber (0.7 µm) filter extracts confirmed the presence of glycosidic extended and di-extended GTGTs (1G-ext-GTGT-0 and 1G-di-ext-GTGT-0) at *m/z* 1553.496 and *m/z* 1623.574 (Fig. 7). Although these lipids were only detected in trace amounts, MS^2^ spectra revealed a fragment ion at *m/z* 1374.414 and *m/z* 1444.499, respectively, reflecting the neutral loss of the glycosidic head group and *m/z* 443.4 representing the extended C_25_-sesterterpanyl side chain. In general, analyses of extracts of glass fiber (0.7 µm) filters and PTFE (0.1 µm) filters resulted in highly similar lipid inventories.

**Fig. 7.**
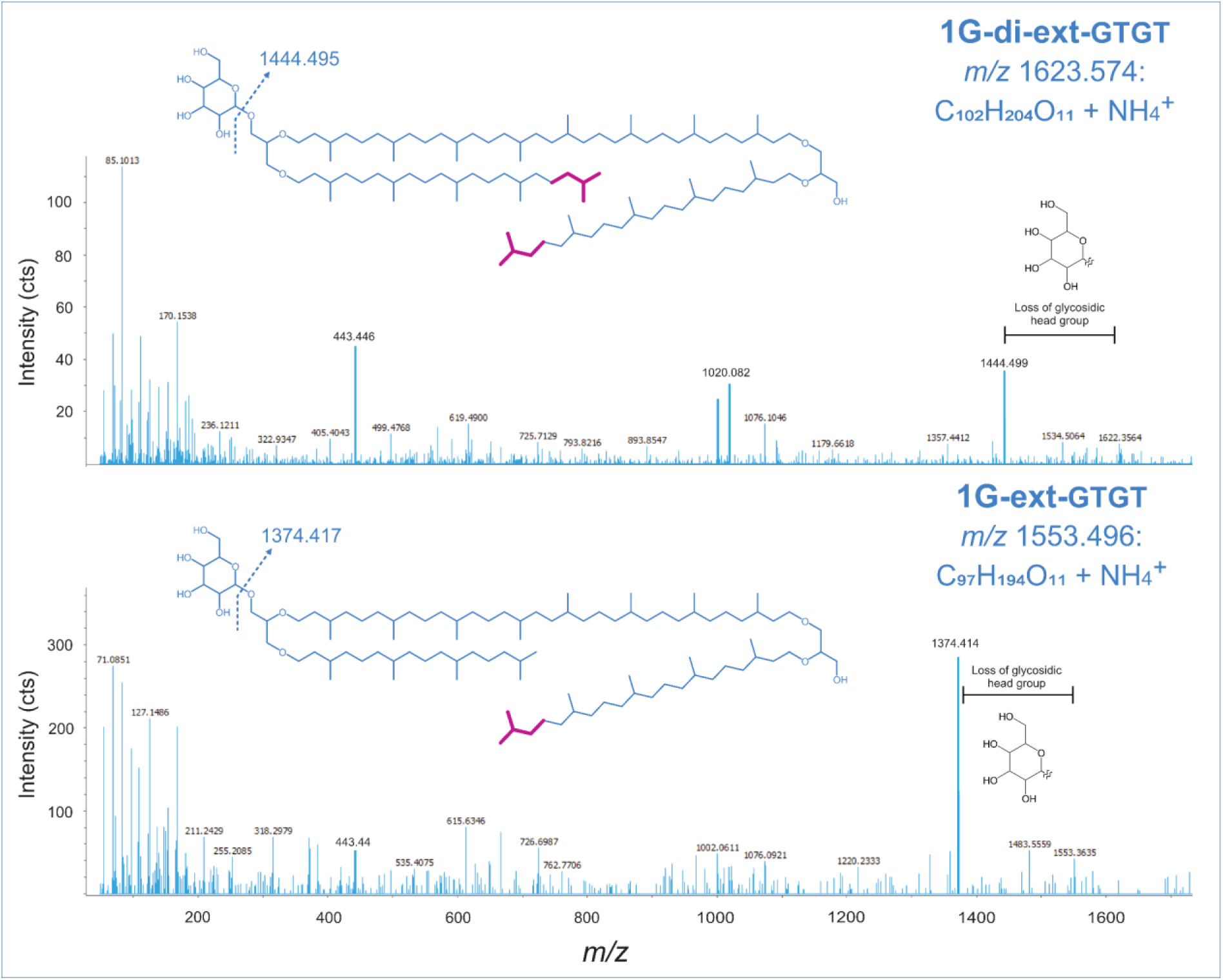
Proposed chemical structures of monoglycosidic extended (C_97_H_194_O_11_) and di-extended GTGT-0 (C_102_H_204_O_11_) found in lipid extracts from the subsurface waters of the Geyser Andernach and corresponding UHPLC-ESI-timsTOF-MS^2^ spectra emphasizing diagnostic MS^2^ fragment ions (method B). The major diagnostic ion arises from the neutral loss of the glycosidic head group. Magenta coloring indicates the additional isoprenoid units characteristic of the extended tetraether lipid structures.

For additional structural information on the C_45_ and C_50_ side chains, GC-MS measurements were attempted. However, the expected C_45_ and C_50_ isoprenoid hydrocarbons, acquired from ether cleavage, reduction, and purification, remained below the detection limit of GC-MS analysis, due to their overall low abundance compared to GDGT-0. Their extended counterparts only represent a minor fraction of the lipidome, accounting for less than 0.5% relative to GDGT-0 (Table S1-2).

### 3.3 Detection of novel extended tetraether lipids beyond the Geyser Andernach

A preliminary re-evaluation of previously conducted lipid measurements from environmental settings, including the Guaymas Basin and the White Oak River, known to host Altiarchaeota of the clade Alti-2 (Bird et al., 2016; Dombrowski et al., 2018; Bornemann et al., 2022), based on metagenomic evidence, revealed a more widespread occurrence of the novel extended tetraether lipids than initially anticipated (Table 1). For instance, ext-GDGT-0 has been detected in the Guaymas Basin sediment, associated with hydrothermally influenced sites. Additionally, di-ext-GDGT-0 occurred, albeit in traces, within the estuarine White Oak River Basin sediment. It is worth noting that the data outside of GA primarily originates from re-inspection of older UHPLC-MS measurements conducted several years ago, when the newly reported extended tetraethers were outside the original analytical focus. Thus, the detection of extended tetraether lipids is qualitative rather than quantitative and should be regarded only as an indication of the occurrence of these lipids at these sites, rather than as an estimate of abundance.

**Table 1.** Peak areas in arbitrary units (arb. u.) of GDGT-0 and its extended analogs (ext-GDGT-0 and di-ext-GDGT-0) found in lipid extracts from hydrothermal sediments in the Guaymas Basin and anoxic estuarine sediments of the White Oak River Basin (WOR). Only peaks with an integrated area above 1000 arb. u. were considered. For each sample, the original measurement and sampling information, including ionization mode (electrospray ionization (ESI) or atmospheric pressure chemical ionization (APCI)) and chromatographic separation, are listed.

| Site | Sediment core/depth | Environmental setting | GDGT-0 Peak area (arb. u.) | ext-GDGT-0 Peak area (arb. u.) | di-ext-GDGT Peak area (arb. u.) | Extended vs typical GDGT-0 % | Ionization mode | LC-separation method | Measurement date | Cruise/Sampling date |
| --- | --- | --- | --- | --- | --- | --- | --- | --- | --- | --- |
| Guaymas Basin | 4568-1_12cm | oil-impregnated, hydrothermal, hot sediment | 1005990 | 1722 | n.d | 0.17 | APCI | normal-phase (Becker et al., 2013) | 03.02.2013 | RV Atlantis cruise AT15-56, 2009 |
| Guaymas Basin | 4484_6B_20-22cm | hot, hydrothermal sediment (Mat Mound) | 6408343 | 38875 | n.d | 0.61 | ESI | reversed-phase (Wörmer et al., 2013) | 07.10.2025 | RV Atlantis cruise AT15-40, 2008 |
| Guaymas Basin | 4484_6B_14-16cm | hot, hydrothermal sediment (Mat Mound) | 14642435 | 107333 | n.d | 0.73 | ESI | reversed-phase (Wörmer et al., 2013) | 07.10.2025 | RV Atlantis cruise AT15-40, 2008 |
| Guaymas Basin | 4484_6B_10-12cm | hot, hydrothermal sediment (Mat Mound) | 14389740 | 69681 | n.d | 0.48 | ESI | reversed-phase (Wörmer et al., 2013) | 07.10.2025 | RV Atlantis cruise AT15-40, 2008 |
| Guaymas Basin | 4484_6B_6-8cm | hot, hydrothermal sediment (Mat Mound) | 9998597 | 23102 | n.d | 0.23 | ESI | reversed-phase (Wörmer et al., 2013) | 07.10.2025 | RV Atlantis cruise AT15-40, 2008 |
| Guaymas Basin | 4484_6B_0-2cm | hot, hydrothermal sediment (Mat Mound) | 17750026 | 11481 | n.d | 0.06 | ESI | reversed-phase (Wörmer et al., 2013) | 07.10.2025 | RV Atlantis cruise AT15-40, 2008 |
| Guaymas Basin | 4869_3_8-10cm | hot, hydrothermal sediment (Ultra Mound) | 10170058 | 80404 | n.d | 0.79 | ESI | reversed-phase (Wörmer et al., 2013) | 28.01.2020 | RV Atlantis cruise AT37-06, 2016 |
| White Oak River Basin | WOR 4-6cm | anoxic, estuarine sediment | 1231649 | n.d | 1059 | 0.09 | APCI | normal-phase (Becker et al., 2013) | 22.01.2016 | 2010 |
| White Oak River Basin | WOR 12-14cm | anoxic, estuarine sediment | 1625887 | n.d | 1390 | 0.09 | APCI | normal-phase (Becker et al., 2013) | 22.01.2016 | 2010 |
| White Oak River Basin | WOR 14-16cm | anoxic, estuarine sediment | 1325301 | n.d | 1298 | 0.10 | APCI | normal-phase (Becker et al., 2013) | 22.01.2016 | 2010 |

### 3.4 Genomic evidence for GDGT synthesis by *Ca.* Altiarchaeum and a distinct evolutionary history of the corresponding gene across the *Ca.* Altiarchaeota phylum

We identified a single *tes* gene homolog in the Geyser Andernach metagenome assembly using DIAMOND BLASTp (identity >50%, coverage >50% and e-value <1e^-5^). This gene was assigned to a previously published *Ca.* Altiarchaeum MAG (accession GCA_018260755.1; GTDB, Release R10-RS226) from the Alti-1 clade (Altiarchaeales), which was recovered from the same metagenome (Bornemann et al., 2022). By contrast, homologs of the GDGT ring synthase genes *grsA* and *grsB* were not detected in the GA metagenome assembly using the same approach.

To recover the evolutionary relationships of the *tes* genes in the *Ca.* Altiarchaeota phylum, we reconstructed a maximum-likelihood phylogenetic tree including the tes gene from this study and reference tes sequences from a previously published dataset (Zeng et al., 2022). Additionally, we included *tes* gene sequences from Alti-2 clade of the Altiarchaeota obtained from Guaymas Basin samples (GCA_003663045.1; GCA_003663165.1; GCA_002254565.1) and White Oak River Basin samples (GCA_001723855.1), in which extended tetraether lipids were detected (Table 1). The resulting tree suggests that the altiarchaeotal *tes* genes may have two distinct evolutionary origins: the *tes* sequence representative from the Geyser Andernach Alti-1 clade clusters with sequences from the *Euryarchaeota* phylum, whereas the Alti-2 clade appears more closely related to sequences from the TACK superphylum (Fig. S4). Additional data will be required to assess potential differences in *tes* genes depending on the types of GDGT modifications observed.

## 4 Discussion

The subsurface waters of the GA harbor a distinct microbial population, where *Ca.* Altiarchaeum represents the sole detected archaeal lineage, recruiting for 42.8% of metagenomic reads (Bornemann et al., 2022). Consequently, we regard *Ca.* Altiarchaeum as the primary and likely exclusive source of the archaeal lipids detected in the geyser waters. The GA archaeal lipidome (Fig. 1) is strikingly consistent with the initial lipid characterization of *Ca.* Altiarchaeum (SM1-MSI strain, Mühlbacher Schwefelquelle) by Probst et al., (2014), who reported a strong dominance of glycosidic ARs (total 91%) accompanied by minor contributions of glycosidic ext-ARs. Contrary to our observations in the GA, neither GDGTs nor extended GDGTs or GTGTs have been reported for a cold sulfidic spring near Regensburg by Probst et al., (2014).

The extension of isoprenoid chains in archaeal membrane lipids has already been well documented in archaeal diether lipids (De Rosa et al., 1982; Vandier et al., 2021). In contrast to the ubiquitously distributed archaeol (Tourte et al., 2020), extended archaeol is typically much more constrained and considered a lipid biomarker for halophilic *Euryarchaeota*, such as *Halobacteriales*, *Haloferacales*, and *Natrialbales* (Dawson et al., 2012) and is often used as an indicator for hypersaline or evaporitic environments (Birgel et al., 2014). However, we note that there are known exceptions, where ext-ARs have been identified in non-halophilic methanogens: *Methanomassilicoccus luminyensis* (Becker et al., 2016) or *Methanosarcina barkerii* (De Rosa et al., 1986), marine lithological sequences with no evidence of hypersalinity (Natalicchio et al., 2017) and multiple reports related to *Ca*. Altiarchaeum (Probst et al., 2014; Perras et al., 2015; Probst et al., 2020), including the cold-water GA, investigated in this study. The most prominent hypothesis explaining the occurrence of ext-AR in *Haloarchaea* relates to their role in mitigating strong osmotic stress. The incorporation of longer, asymmetric C_20_-C_25_ AR in a potentially interlocking structure increases membrane thickness and reduces passive ion leakage (De Rosa et al., 1982), which in turn helps to maintain the proton motive force under extreme salinity. Since the GA water is best described as brackish, we hypothesize that its nutrient- and energy-limited character, rather than salinity, act as a comparable stressor, driving these physiological adaptations (Bornemann et al., 2022).

A novel finding of our study was the detection of GDGT-0 and its intact counterpart 1G-GDGT-0, marking the first report of tetraether lipids attributable to the dominating microbial population of the GA, *Ca.* Altiarchaeum (Bornemann et al., 2022). Interestingly, a recent genomic survey using Basic Local Alignment Search Tool (BLASTp) already suggested that Altiarchaeota have the potential genetic capacity for GDGT biosynthesis, as homologs of the tetraether synthase (*tes*) have been identified within their genome (Zeng et al., 2022). This radical S-adenosylmethionine (SAM) enzyme plays a critical role in the formation of GDGTs by catalyzing the coupling of two isoprenoidal diether lipids, primarily archaeols, under the formation of glycerol trialkyl glycerol tetraether (GTGTs) as intermediates (Koga and Morii, 2007; Matsumi et al., 2011; Straub et al., 2018; Lloyd et al., 2022; Zeng et al., 2022). This biosynthetic potential is corroborated by the detection of a *tes* homolog, with a strong match to reference sequences, in a *Ca.* Altiarchaeum (Alti-1 clade) MAG recovered from GA (Bornemann et al., 2022) (cf. section 3.4).

Beyond GDGT-0, the GA lipidome also features novel GDGTs and GTGTs comprising up to two additional isoprenoid units similar to those described in the archaeal C_20_-C_25_ diethers. Given that Altiarchaeales are the dominant prokaryotic and sole archaeal group within the geyser subsurface (Bornemann et al., 2022), and appear to be the primary GDGT-producing archaea in this setting, it is most probable that these structurally related tetraether lipids originate from the same source. It is possible that these modified extended GDGTs and GTGTs reflect an expanded versatility in the tetraether synthesis in Altiarchaeales, potentially as an adaptation to the unique physicochemical conditions in the deep subsurface environment.

Surprisingly, ext-GTGT and di-ext-GTGT were more abundant than their dialkyl GDGT counterparts. This pattern is also reflected in their glycosylated derivatives, as only 1G-ext-GTGT-0 and 1G-di-ext-GTGT-0 were detected in trace amounts (Fig. 1), while intact ext-GDGTs remained below the detection limit. GTGTs are considered a transient intermediate in GDGT biosynthesis, consistent with their typically low abundance in both environmental samples and cultures (Liu et al., 2012b; Bauersachs et al., 2015; Tourte et al., 2022). This may indicate that Tes processes the ext-AR substrates less efficiently. The larger C_25_-chain potentially slows the second enzymatic condensation step, creating a kinetic bottleneck and, in turn, causing the accumulation of the extended GTGT intermediates. However, this interpretation is based on observations from a single site, and more targeted studies are needed to determine whether there are differences in the biosynthesis of regular and extended GDGTs.

Common adaptation strategies to environmental stressors, such as temperature, often involve incorporating up to eight cyclopentyl rings into the isoprenoid chains (De Rosa and Gambacorta, 1988; Lai et al., 2008; Boyd et al., 2011). The insertion of ring moieties increases packing density and rigidifies the lipid membrane, enabling archaea to maintain cell integrity and functionality and prevent ion leakage under extreme temperatures or pH (De Rosa et al., 1980; Gliozzi et al., 1983; Chiu et al., 2023). Considering the absence of GrsA and GrsB homologs, required for the formation of cyclopentane rings (Zeng et al., 2019) in the GA metagenome, it is probable that members of this phylum do not have the biosynthetic capability to produce cyclized GDGTs. Although additional UHPLC-APCI-MS analysis (Table S3) revealed traces of cyclopentyl-bearing GDGTs, the absence of GrsA and GrsB homologs makes it unlikely that Altiarchaeales are the source of these lipids. It is conceivable that another, low abundance archaeal taxon not captured in the metagenome may contribute those trace amounts to the lipid inventory. Nevertheless, the isoprenoid extension of tetraether lipids is likely attributable to the predominant Altiarchaeales and may represent a novel adaptation strategy to prevent ion leakage in the nutrient-depleted cold-water aquifer (Bornemann et al., 2022).

Overall, the close association of these unusual extended tetraether lipids and *Ca*. Altiarchaeales hints that they may serve as a chemotaxonomic indicator for this lineage. However, additional comprehensive lipidomic surveys from sites known to host Altiarchaeota, as well as reference sites, are needed to evaluate their true biomarker potential. Alternatively, the occurrence of extended GDGTs may reflect a more general environmental adaptation to extreme subsurface habitats, particularly in archaea, which lack the capacity for GDGT cyclization. To explore whether these novel GDGTs occur beyond the GA subsurface, we performed an initial re-inspection of preexisting lipid measurements from environments reported to host Altiarchaeota of either Alti-1 clade or Alti-2 clade (Bird et al., 2016), and detected traces of extended tetraether lipids (Table 1). Most prominent was the detection of ext-GDGT-0 in the hydrothermally influenced deep-sea sediment from the Guaymas Basin and di-ext-GDGT-0 in anoxic, estuarine sediment from the White Oak River Basin. Both are sites where Altiarchaeota, particularly of the Alti-2 clade, which also carry the *tes* gene required for tetraether synthesis (cf. section 3.4), have been reported (Bird et al., 2016; Dombrowski et al., 2018; Bornemann et al., 2022). However, in contrast to GA, where Altiarchaeota dominate, the Guaymas Basin and the White Oak River Basin host considerably more diverse archaeal communities, resulting in a less straightforward source attribution. Although these extended lipids cannot be confidently linked to Altiarchaeota at those sites, their detection still provides a first indication that extended tetraether lipids occur in a wide range of environmental settings, including subsurface aquifers, hydrothermal marine sediment, and anoxic estuarine sediment. Due to their relatively low abundance compared to typical GDGTs (Fig. 1 and Table 1), they may have escaped detection in previous studies and could be more widely distributed than currently recognized. Future lipidomic investigations employing high-sensitivity analytical approaches, ideally combined with genomic data, will be needed to resolve the ecological significance as well as their microbial origin across diverse environmental settings. Additionally, molecular dynamics simulations and membrane modeling of regular GDGTs versus extended GDGTs could provide valuable insight into their biophysical properties and their effects on membrane stability and permeability, particularly in extreme environments.

## 5 Conclusion

Our findings add a novel group of isoprenoidal tetraethers to the steadily growing catalogue of archaeal membrane lipids, further underscoring their remarkable structural diversity. The newly detected extended and di-extended GTGTs and GDGTs are characterized by the addition of isoprenoid units, analogous to those described in ext-AR, while retaining one conventional biphytanyl motif, ultimately resulting in an asymmetrical structure. The dominance of Altiarchaeota in the CO_2_-rich, nutrient-depleted subsurface waters of the GA, combined with the only Tes homolog being affiliated with *Ca.* Altiarchaeum strongly implies this lineage as their most probable source. We hypothesize that, at this site, *Ca*. Altiarchaeum employs the extension of isoprenoidal side chains as an alternative strategy for adapting membrane permeability and energy conservation under nutrient- and energy-limited conditions.

## 6 Acknowledgements

We thank Lukas Dirksen for his assistance with instrumental analyses. We also thank the “Archean-Park” team members from the University of Duisburg-Essen for providing their filtration apparatus.

## Notes

### Competing Interest Statement

The authors have declared no competing interest.

### Funding

- European Research Council: 101118631

