## Supplementary material for "Novel Extended Tetraether Lipids Found in a High-CO_2_ Geyser"

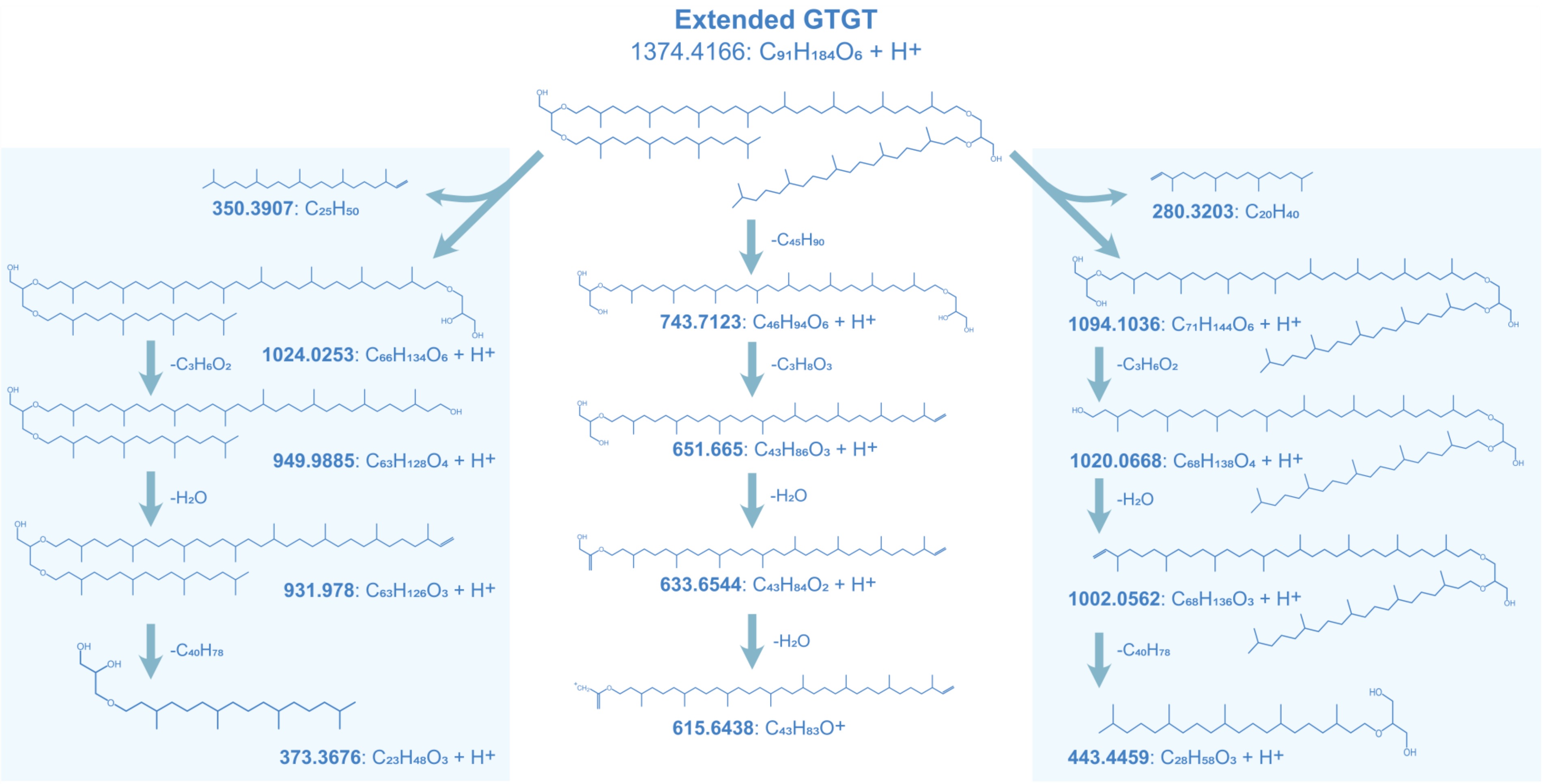


**Fig. S1.** Proposed fragmentation of ext-GTGT-0 $[M+H]^+$ based on $MS^2$ spectra acquired via APCI-QTOF-$MS^2$ in positive ionization mode.

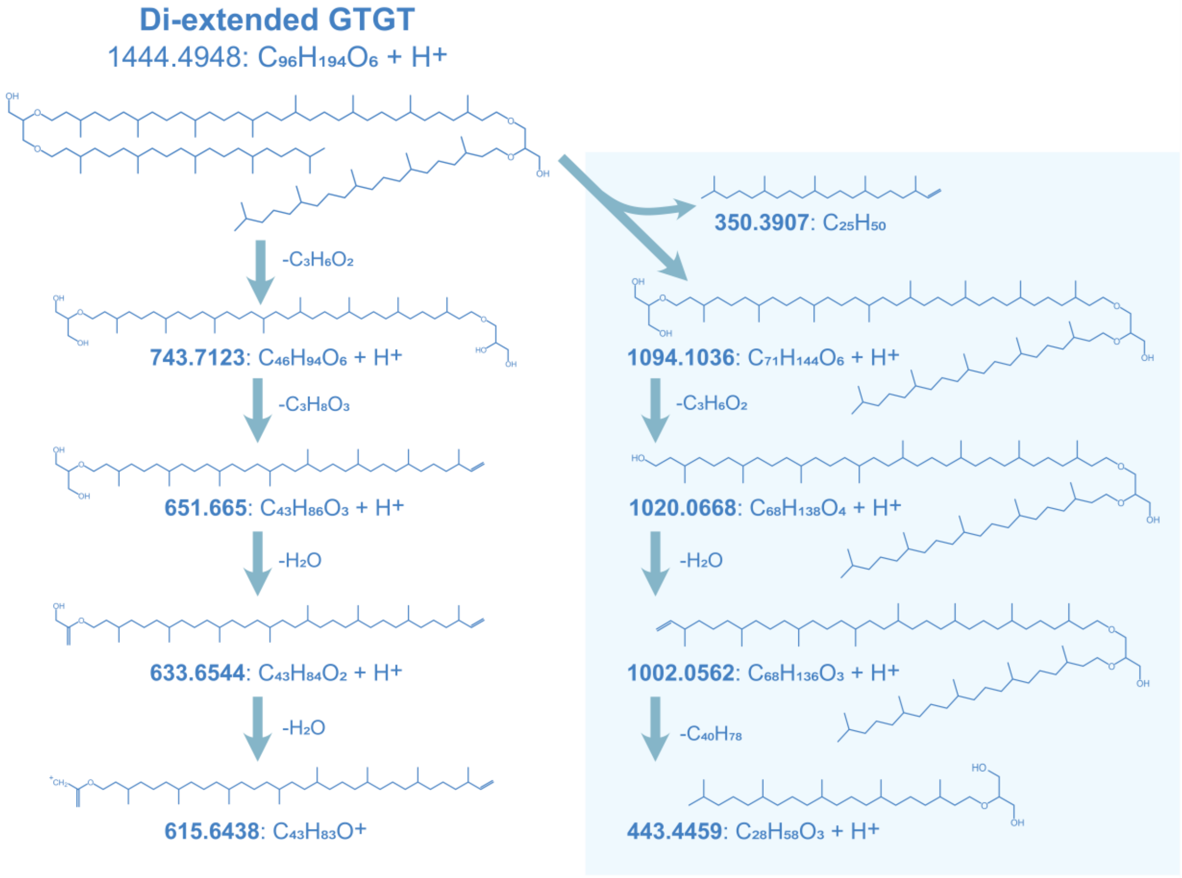


**Fig. S2.** Proposed fragmentation of di-ext-GTGT-0 $[M+H]^+$ based on $MS^2$ spectra acquired via APCI-QTOF-$MS^2$ in positive ionization mode.

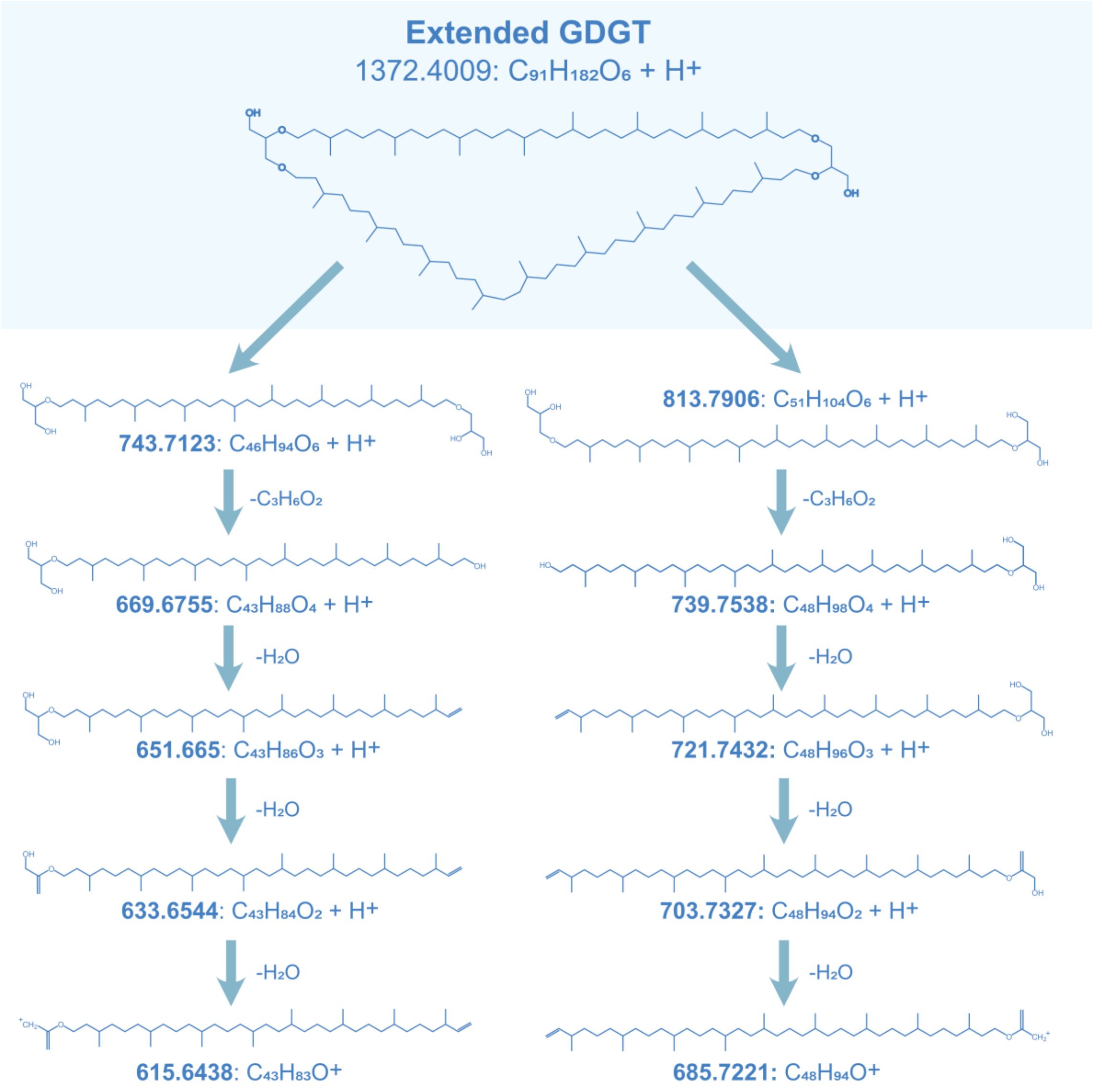


**Fig. S3.** Proposed fragmentation of ext-GDGT-0 $[M+H]^+$ based on $MS^2$ spectra acquired via APCI-QTOF-$MS^2$ in positive ionization mode.

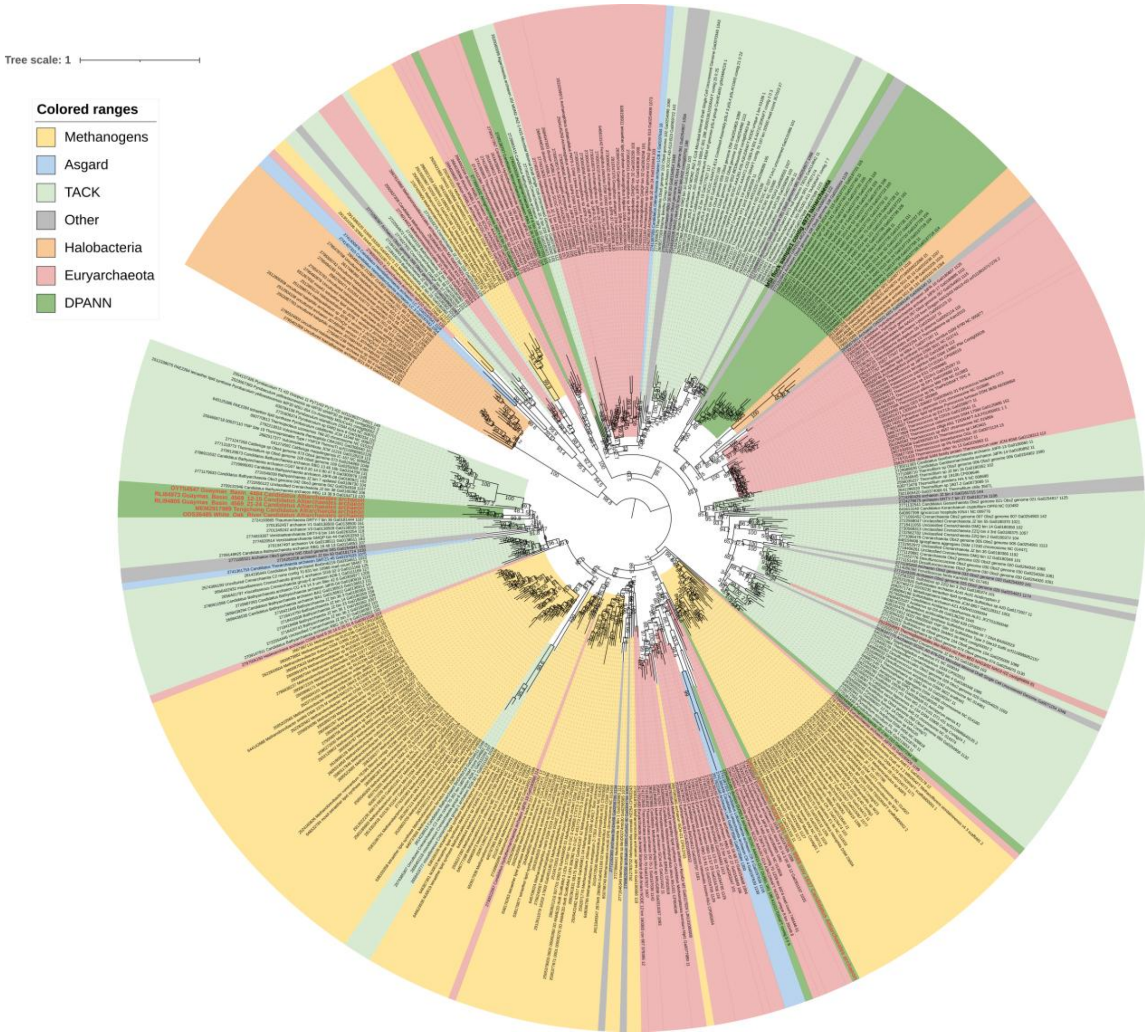


**Fig. S4.** Maximum-likelihood phylogenetic tree about the evolutionary relationships of the *tes* genes, including references from Zeng et al. 2022. Additional archaeal sequences from the Geyser Andernach metagenome of this study, Guaymas Basin (Core 4484 and 4569) and White Oak River Basin are marked in red and resemble sites, in which the both, the *tes* gene and extended tetraether lipids were found.

**Table S1.** Overview of **all identified archaeal lipids** identified in the erupting water of the Geyser Andernach via UHPLC-ESI-timsTOF-MS. The reported parameters include the chemical formula, monoisotopic mass, three most abundant adducts, retention time (using method A) as well as peak area (arbitrary units), relative abundance, concentration (ng per TLE) and inverse reduced ion mobility values of the $[M+NH_4]^+$ adducts.

| Lipid | Chem. formula | Monoisotopic mass | *m/z* Adducts $[M+H]^+$; $[M+NH_4]^+$; $[M+Na]^+$ | RT [min] | Peak Area [arb. u.] | Rel. Abundance [%] | Conc. [ng/TLE] | Mobility $1/K_0$ [Vs cm$^{-2}$] |
|---|---|---|---|---|---|---|---|---|
| **1G-AR** | $C_{49}H_{98}O_8$ | 814.726 | 815.733; 832.760; 837.715 | 16.8 | 7126755 | 15 | 6.20 | 1.53 ± 0.02 |
| **2G-AR** | $C_{55}H_{108}O_{13}$ | 976.778 | 977.786; 994.8128; 999.768 | 15.6 | 17161294 | 35 | 14.93 | 1.65 ± 0.02 |
| **3G-AR** | $C_{61}H_{118}O_{18}$ | 1138.381 | 1139.839; 1156.866; 1161.821 | 14.8 | 1725292 | 4 | 1.50 | 1.74 ± 0.02 |
| **AR** | $C_{43}H_{88}O_3$ | 652.673 | 653.681; 670.707; 675.663 | 18.6 | 4354194 | 2 | 0.72 | 1.38 ± 0.02 |
| **1G-GDGT-0** | $C_{92}H_{182}O_{11}$ | 1463.368 | 1464.375; 1481.402; 1486.357 | 23.5 | 2432768 | 9 | 3.99 | 2.06 ± 0.02 |
| **GDGT-0** | $C_{86}H_{172}O_6$ | 1301.315 | 1302.323; 1319.349; 1324.305 | 24.4 | 53530108 | 25 | 10.69 | 1.98 ± 0.02 |
| **Extended Lipids** | - | - | - | - | - | 10 | - | - |

**Table S2.** Overview of **extended archaeal lipids** identified in the erupting water of the Geyser Andernach via UHPLC-ESI-timsTOF-MS. The reported parameters include the chemical formula, monoisotopic mass, three most abundant adducts, retention time (using method A) as well as peak area (arbitrary units), relative abundance, concentration (ng per TLE) and inverse reduced ion mobility values of the $[M+NH_4]^+$ adducts.

| Lipid | Chem. formula | Monoisotopic mass | Adducts $[M+H]^+$; $[M+NH_4]^+$; $[M+Na]^+$ | RT [min] | Peak Area [arb. u.] | Rel. Abundance [%] | Conc. [ng/TLE] | Mobility $1/K_0$ [Vs cm$^{-2}$] |
|---|---|---|---|---|---|---|---|---|
| **1G-ext-AR** | $C_{54}H_{108}O_8$ | 884.804 | 885.812; 902.838; 907.794 | 19.0 | 1511827 | 30 | 1.32 | 1.61 ± 0.02 |
| **2G-ext-AR** | $C_{60}H_{118}O_{13}$ | 1046.857 | 1047.864; 1064.891; 1069.846 | 17.9 | 1320736 | 26 | 1.15 | 1.71 ± 0.02 |
| **3G-ext-AR** | $C_{66}H_{128}O_{18}$ | 1208.909 | 1209.917; 1226.944; 1231.899 | 17.3 | 191498 | 4 | 0.17 | - |
| **ext-AR** | $C_{48}H_{98}O_3$ | 722.751 | 723.759; 740.785; 745.741 | 20.7 | 980529 | 4 | 0.16 | 1.46 ± 0.02 |
| **ext-GTGT-0** | $C_{91}H_{184}O_6$ | 1373.409 | 1374.416; 1391.443; 1396.398 | 25.3 | 3849205 | 18 | 0.77 | 2.09 ± 0.02 |
| **di-ext-GTGT-0** | $C_{96}H_{194}O_6$ | 1443.487 | 1444.495; 1461.521; 1466.477 | 26.3 | 1683153 | 8 | 0.34 | 2.15 ± 0.02 |
| **1G-ext-GTGT-0** | $C_{97}H_{194}O_{11}$ | 1535.461 | 1536.469; 1553.496; 1558.451 | 24.2 | 183218 | 7 | 0.30 | 2.17 ± 0.02 |
| **1G-di-ext-GTGT-0** | $C_{102}H_{204}O_{11}$ | 1605.540 | 1606.548; 1623.574; 1628.529 | 24.9 | 55766 | 2 | 0.09 | 2.23 ± 0.02 |
| **ext-GDGT-0** | $C_{91}H_{182}O_6$ | 1371.393 | 1372.401; 1389.427; 1394.383 | 25.1 | 221317 | 1 | 0.04 | 2.08 ± 0.02 |
| **di-ext-GDGT-0** | $C_{96}H_{192}O_6$ | 1441.472 | 1442.479; 1459.506; 1464.461 | 26.0 | 97050 | 0.4 | 0.02 | 2.14 ± 0.02 |

**Table S3.** Overview of **archaeal core lipids** obtained via UHPLC-APCI-QTOF-MS measurements of erupting water of the Geyser Andernach including their chemical formula, the most abundant adduct, *m/z*, retention time as well as peak area in arbitrary units (arb. u.) and relative abundance.

| Lipid | Chem. Formula | Adducts | *m/z* | RT [min] | Peak Area [arb. u.] | Rel. Abundance [%] |
|---|---|---|---|---|---|---|
| **di-ext-GTGT-0** | $C_{96}H_{194}O_6$ | $[M+H]^+$ | 1444.4948 | 14.4 | 777716 | 3.7 |
| **di-ext-GDGT-0** | $C_{96}H_{192}O_6$ | $[M+H]^+$ | 1442.4792 | 17.4 | 18963 | 0.1 |
| **ext-GTGT-0** | $C_{91}H_{184}O_6$ | $[M+H]^+$ | 1374.4166 | 14.8 | 1689421 | 8.1 |
| **ext-GDGT-0** | $C_{91}H_{182}O_6$ | $[M+H]^+$ | 1372.4009 | 17.9 | 270903 | 1.3 |
| **GDGT-0** | $C_{86}H_{172}O_6$ | $[M+H]^+$ | 1302.3227 | 18.4 | 14473976 | 69.5 |
| **GTGT-0** | $C_{86}H_{174}O_6$ | $[M+H]^+$ | 1304.3383 | 15.2 | 2847365 | 13.7 |
| **GDGT-1** | $C_{86}H_{170}O_6$ | $[M+H]^+$ | 1300.3070 | 18.6 | 254546 | 1.2 |
| **GDGT-1 (Isomer)** | $C_{86}H_{170}O_6$ | $[M+H]^+$ | 1300.3070 | 18.9 | 73570 | 0.4 |
| **GDGT-2** | $C_{86}H_{168}O_6$ | $[M+H]^+$ | 1298.2914 | 19.5 | 82204 | 0.4 |
| **GDGT-3** | $C_{86}H_{166}O_6$ | $[M+H]^+$ | 1296.2757 | 20.2 | 42081 | 0.2 |
| **GDGT-4** | $C_{86}H_{164}O_6$ | $[M+H]^+$ | 1294.2601 | 20.9 | 12988 | 0.1 |
| **GMGT-0** | $C_{86}H_{170}O_6$ | $[M+H]^+$ | 1300.3070 | 22.2 | 219307 | 1.1 |
| **GMGT-1** | $C_{86}H_{168}O_6$ | $[M+H]^+$ | 1298.2914 | 22.8 | 72509 | 0.3 |
